# PHACE: Phylogeny-Aware Co-Evolution

**DOI:** 10.1101/2024.12.19.629429

**Authors:** Nurdan Kuru, Ogun Adebali

## Abstract

The co-evolution trends of amino acids within or between genes offer valuable insights into protein structure and function. Existing tools for uncovering co-evolutionary signals primarily rely on multiple sequence alignments (MSAs), often neglecting considerations of phylogenetic relatedness and shared evolutionary history. Here, we present a novel approach based on the substitution mapping of amino acid changes onto the phylogenetic tree. We categorize amino acids into two groups: ‘tolerable’ and ‘intolerable,’ and assign them to each position based on the position dynamics concerning the observed amino acids. Amino acids deemed ‘tolerable’ are those observed phylogenetically independently and multiple times at a specific position, signifying the position’s tolerance to that alteration. Gaps are regarded as a third character type, and we only consider phylogenetically independent altered gap characters. Our algorithm is based on a tree traversal process through the nodes and computes the total amount of substitution per branch based on the probability differences of two groups of amino acids and gaps between neighboring nodes. We employ an MSA-masking approach to mitigate misleading artifacts from poorly aligned regions. When compared to tools utilizing phylogeny (CAPS and CoMap) and state-of-the-art MSA-based approaches (DCA, GaussDCA, PSICOV, and MIp), our method exhibits significantly superior accuracy in identifying co-evolving position pairs, as measured by statistical metrics including MCC, AUC, and F1 score. PHACE’s success arises from its ability to consider the frequently neglected phylogenetic dependency.

## Introduction

Coevolution refers to the synchronized alterations observed in pairs of organisms or biomolecules, typically aimed at preserving or enhancing the functional relationships between them (De Juan et al. 2013). While it occurs across various levels, such as among species and organisms, it is particularly evident at the molecular level between interacting protein positions (Dutheil 2012). The literature has shown significant interest in detecting molecular coevolution and understanding the trends of coevolution among protein positions, as it offers vital insights into protein structure and function. Notably, cutting-edge methodologies like AlphaFold (Jumper et al. 2021) and RoseTTAFold (Baek et al. 2021) leverage covariation as a crucial input feature, underscoring its importance in modern protein structure prediction.

Co-evolution trends between amino acid positions can be detected using various approaches based on correlated changes. Many approaches based on multiple sequence alignments (MSAs) are presented in the literature to detect co-evolution, such as the state-of-the-art tools, DCA (Morcos et al. 2011), GaussDCA (Baldassi et al. 2014), mutual information (MIp) (Dunn et al. 2008), and PSICOV (Jones et al. 2012b). However, coevolution is not the sole source of information resulting in correlated amino acid presences between protein positions (Dutheil 2012). Thus, it is essential to discriminate the actual co-evolution signal from other sources of correlated changes, where a primary false signal is known to be caused by phylogenetic relatedness (Dutheil 2012). Additionally, methods scoring co-evolution based on the covariation of positions are known to fail in discriminating positions differentiating in evolutionary scenarios (Talavera et al. 2015). Talavera et al. demonstrated the indistinguishability of co-evolutionary scenarios from non-coevolving scenarios based solely on covariation, highlighting its limitations as a measure of co-evolution.

Fig 1 illustrates our rationale, demonstrating that even if position pairs exhibit identical amino acid frequencies in the MSA, the number of correlated changes can vary significantly depending on the topology of the phylogenetic tree (Fig 1A). In the first tree, four correlated changes are observed as evolutionary independent, while all four substitutions on the second tree are phylogenetically dependent and they occurred due to a single amino acid alteration. In other words, these two scenarios are equivalent in terms of MSA-based scoring of the co-evolution signal; however, ignoring common evolutionary history results in a false co-evolution signal and overcounting the effect of one single amino acid alteration as if it occurred four times independently. Our study aims to address this challenge by accurately scoring genuine co-evolution signals resulting from correlated evolutionary variation while excluding phylogenetic relatedness.

**Figure 1.**
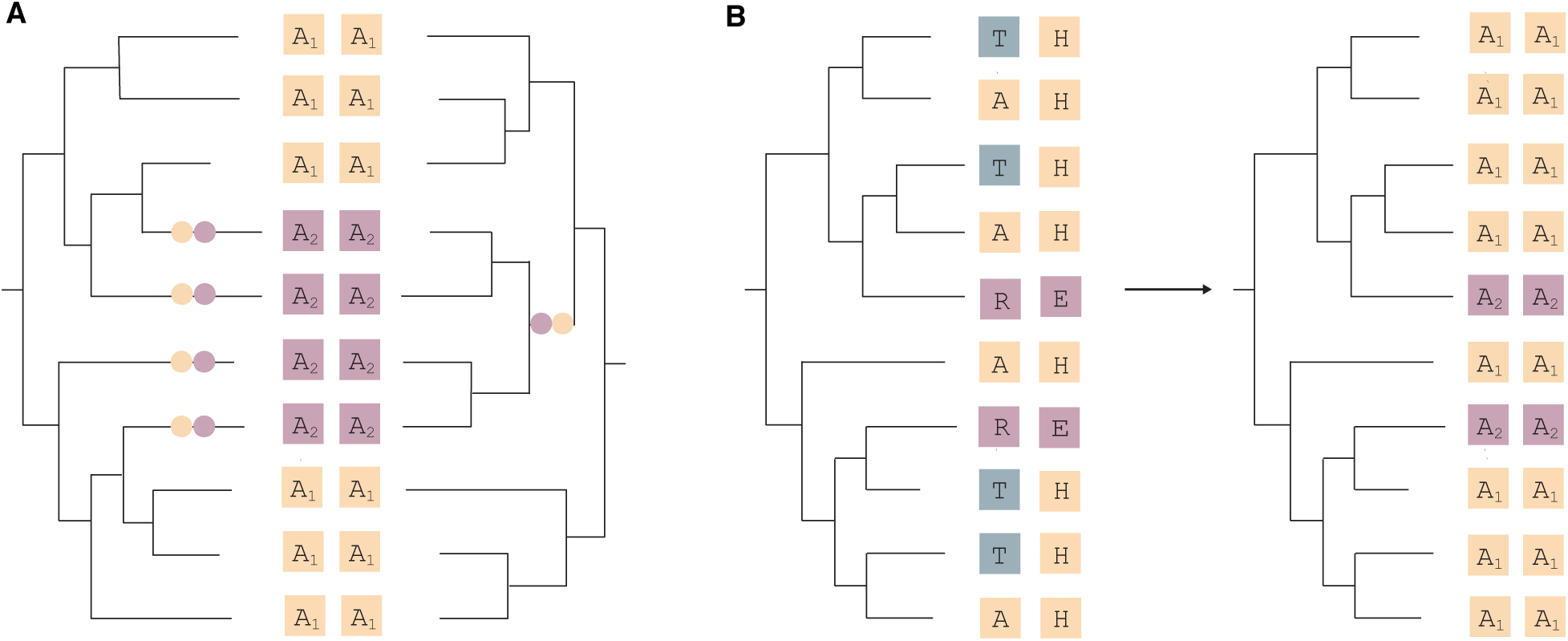
Rationale of PHACE. **A.** Importance of phylogenetic information in co-evolution: Despite identical multiple sequence alignments (MSAs) for position pairs, phylogenetic analysis reveals differing correlated changes. **B.** Clustering amino acids by tolerability: Despite unclear co-evolution signals initially, repeated observation of A’s and T’s at a position suggests tolerance to both amino acids which are represented as A_1_ in the updated MSA, while other amino acids are grouped as A_2_., This process alters the co-evolution signal in the updated MSA.

We emphasize another problem that disrupts the co-evolution signal: high variability in aligned positions. The variability in protein positions due to amino acid changes complicates our understanding of evolution and makes scoring parallel changes difficult. Certain positions can tolerate amino acid changes, thus contributing to sequence divergence without affecting protein function. We have noticed that scoring this kind of substitution similarly to an intolerable substitution in the relevant position can hinder the co-evolution signal. An illustrative example is provided in Fig 1B. Despite the uncertain co-evolution signal in the original MSA, both Alanines (As) and Threonines (Ts) are observed phylogenetically independently and repeatedly, highlighting the position’s tolerance to both A and T. We incorporate this tolerance into our framework and group amino acids into two clusters based on whether they are tolerated or not as A_1_ and A_2_, respectively. Since As and Ts are labelled as ‘tolerated’ in our framework, we behave them the same by grouping them as A_1_. The remaining amino acids which are the ones we use to score co-evolution signal are clustered as A_2_. As illustrated in the updated MSA in Fig 1B, this approach reveals the co-evolution signal concealed in the original MSA.

Several attempts have been made in the literature to solve the first problem: to separate co-evolution signals from phylogenetic relatedness by incorporating phylogenetic trees. CAPS (Fares and McNally 2006) and CoMap (Dutheil and Galtier 2007), in particular, leverage both phylogenetic trees and ancestral sequence reconstruction in their scoring computations. CAPS detects coevolving amino acid sites by measuring the correlation of evolutionary rate variability between sites, corrected for divergence time. In the initial version of CAPS, phylogenetic trees were used solely for correction; however, in CAPS v2, which is the version considered here, the approach is based on substitution mapping of amino acid changes onto the phylogenetic tree. CoMap is another tree-based approach that benefits from substitution mapping. Although both approaches use trees and ancestral sequence reconstruction (ASR), they assign a single amino acid to internal nodes, rely on a simple correlation matrix, and ignore position dynamics. In our previous work (Kuru et al. 2022b), we introduced PHACT, a novel phylogeny-based approach for predicting the functional consequences of missense mutations. PHACT assesses amino acid substitutions using phylogenetic trees and the probabilities derived from ancestral reconstructions. It employs a tree traversal method that tracks positive probability differences between adjacent nodes, beginning from the leaf of the query sequence. PHACT effectively utilizes phylogenetically independent events to predict the pathogenicity of missense mutations through this tree traversal process. Given its success in scoring phylogenetically independent events by accurately eliminating the effect of shared evolutionary history, we have developed PHACE, a novel phylogeny-aware co-evolution algorithm.

The PHACE method aims to detect parallel substitutions between pairs of positions by leveraging phylogenetically independent events. The outline of the PHACE algorithm is illustrated in Fig 2. We aim to eliminate a significant source of correlated changes, irrelevant to co-evolution, due to phylogenetic relatedness. We achieve this by examining the differences in probability distributions between neighboring nodes and calculating the total amount of positive probability differences. This total corresponds to the number of phylogenetically independent amino acid alterations per branch. The probability distribution of amino acids per internal node is obtained at the ASR. Total change per branch is calculated to measure co-evolutionary patterns. However, as mentioned earlier, certain positions can tolerate amino acid changes without affecting protein function. To mitigate this problem and ensure an accurate assessment of position diversity, we utilized a reconstructed a version of an MSA derived from the original MSA (MSA_1_ in Fig 2A). This refined MSA categorizes amino acids into three groups: tolerable, intolerable, and gap characters. “*A*_1_”, “*A*_2_” and “−” characters are used for tolerable, intolerable amino acids and gaps, respectively. The position tolerability to any amino acid is obtained based on phylogenetically independent amino acid alterations.

**Figure 2.**
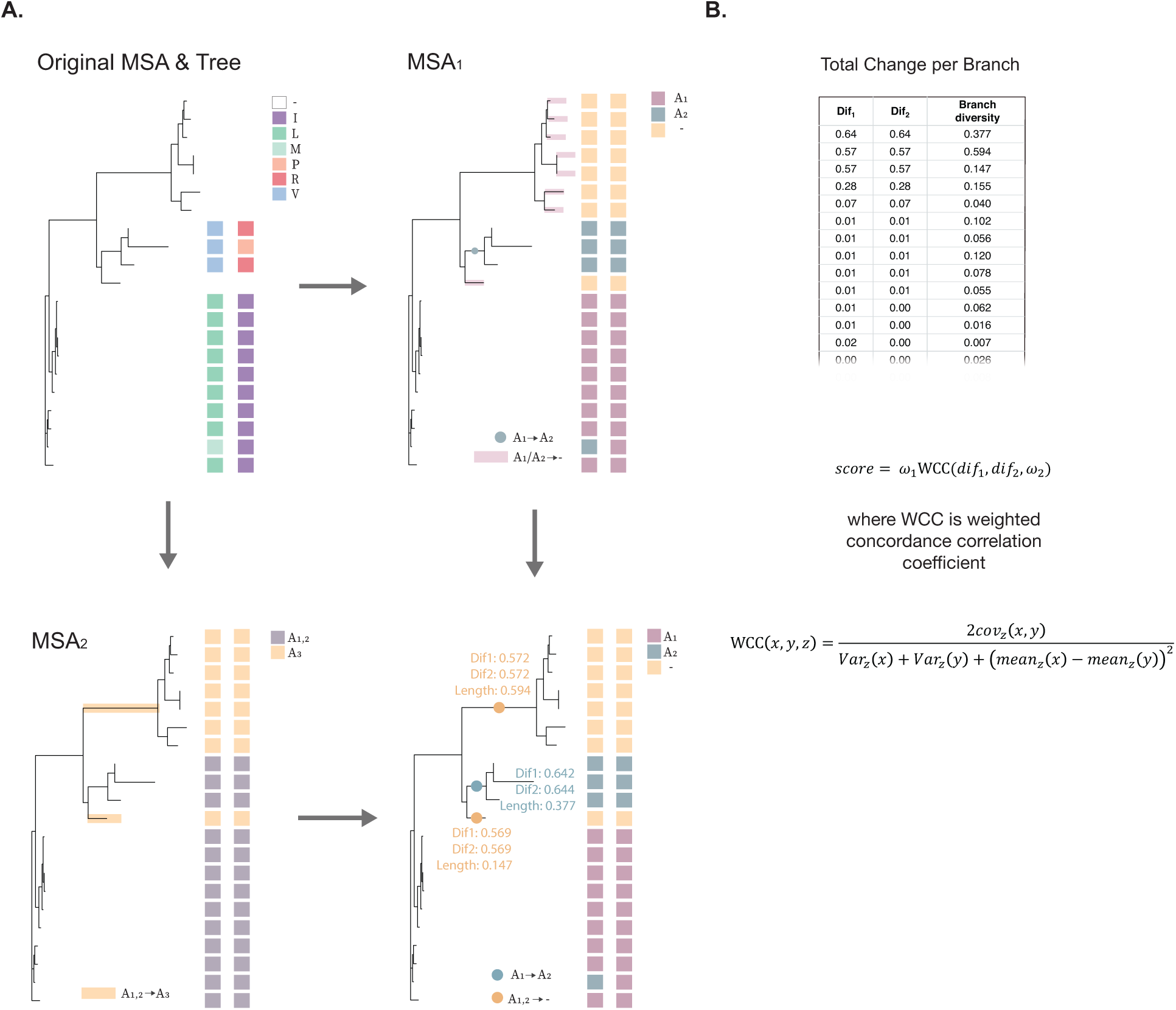
PHACE Algorithm Overview. **A.** PHACE utilizes the original MSA and ML phylogenetic tree to cluster amino acids into “tolerable” and “intolerable” groups, resulting in MSA_1_. To address issues with gapped leaves and obtain accurate co-evolution signals, MSA_2_ is created to distinguish amino acids from gaps. This information is used to update substitution rates per branch from MSA_1_. The final MSA is created by integrating MSA1 and MSA2, considering phylogenetically independent changes. B. The final MSA is then used to construct a matrix detailing changes per branch per position and branch diversity. PHACE score is calculated using a weighted concordance correlation coefficient. (Pos. 126-130, spatial distance: 6.54 Å)

Although we successfully eliminate the correlated patterns resulting from shared evolutionary history and consider position diversity, deciding how to treat gaps is important. In the existing literature, widely-used tools such as DCA, GaussDCA, PSICOV, and Mutual Information treat gaps as the 21st character. However, most tools overlook gaps in sequence reconstruction in the ASR framework. However, evolutionary-independent multiple observations of indels suggest a potential co-evolved site. Ignoring gaps and treating them as the 21^st^ amino acid limits the sensitivity and specificity of identifying the co-evolving sites, respectively. To address these limitations, we introduce a second version of MSA consisting of only two characters: one for all amino acids and one for gaps (MSA_2_ on Fig 2A). By applying classical ASR algorithms to this simplified MSA, we pinpoint the occurrence of phylogenetically independent gap alterations, which correspond to the branches where the probability of the character assigned to the gap increases. As shown in the final tree in Fig 2A, we consider only phylogenetically independent amino acid and gap alterations and eliminate the effect of shared evolutionary history from our co-evolution score with this approach.

By integrating information from both versions of MSAs and their associated ASR probabilities, we generate a matrix with dimensions corresponding to the number of branches by 2, as shown in Fig 2B. Each row in this matrix represents the total amount of phylogenetically independent amino acid alterations. The PHACE score is obtained by utilizing the weighted concordance correlation coefficient (WCCC) over the matrix of total changes per branch and branch diversity, with the latter serving as the weight. Branch diversity helps identify whether variability at a branch is widespread or specific to certain positions. By distinguishing generally diverse branches from those with localized changes, we can more reliably attribute the observed changes to co-evolution rather than to intrinsic variability. The branch diversity is computed by considering the total amount of change for each branch across all positions. Although the branch length of the phylogenetic tree could potentially be used for this purpose, it is noteworthy that, since the gap character is not employed in tree construction and ASR steps, the branch lengths do not accurately represent the overall changeability of the corresponding branch. Hence, we have determined another score that better represents this information.

In our experiments aimed at discriminating between close and distant positions on the 3D structure of human proteins, PHACE demonstrated significantly superior performance across various statistical measures compared to MSA-based tools (DCA, GaussDCA, PSICOV, and MIp) and phylogeny-based approaches (CAPS and CoMap).

## Results

To evaluate the performance of the PHACE algorithm, we utilized protein 3D structures and limited our interest to the proteins with experimentally determined structures. The criteria for determining the protein set are detailed in the Materials and Methods section. Similar to the previous studies, we considered two positions are ‘in contact’ if their Cβ-Cβ distance is less than 8 angstroms (Å) (Morcos et al. 2011; Jones et al. 2012a; Baldassi et al. 2014). Thus, following the literature, we infer that two positions are co-evolving if they are proximate in the 3D structure. While an 8 Å threshold is commonly accepted for defining positions in contact, some studies suggest using distances up to 12 Å. In our analysis, we employed two different strategies for defining non-coevolving position pairs.

First, we used a threshold of 16 Å to classify non-coevolving pairs. This threshold was chosen based on the structural properties of proteins, such as the typical spacing observed within regular secondary structural elements like alpha helices and beta strands. In alpha helices, which have about 3.6 amino acids per turn, each residue contributes to a helical rise of approximately 1.5 Å. Similarly, amino acids in beta strands are spaced roughly 3.5 Å apart along the strand. By setting a 16 Å threshold, we ensured that position pairs separated by distances greater than the spacing within these motifs were categorized as non-coevolving, helping to minimize false positive co-evolution signals. In this approach, positions at least 16 Å apart in the 3D structure were labeled as non-coevolving.

Second, for ROC curve comparisons with tools such as DCA, GaussDCA, PSICOV, and MIp, we implemented a strategy where non-coevolving pairs were chosen by sorting distances from the farthest to the closest, up to the 8 Å threshold, to match the number of co-evolving pairs. This balanced selection approach addressed potential biases in the data set, which can impact metrics like AUC that are sensitive to imbalances. Using these two complementary strategies, we aimed to maximize the reliability of our assessments and ensure consistency across different analyses.

As benchmark tools report results in various formats, we selected statistical measures based on their respective outputs. CAPS and CoMap exclusively report co-evolving position pairs, while DCA and GaussDCA provide scores for all pairs. Consequently, we evaluate CAPS and CoMap using Matthews correlation coefficient (MCC) and F1 scores, whereas the area under the ROC curve (AUC) is utilized for comparing performance with DCA, GaussDCA, PSICOV and MIp since they report predictions for all (almost) position pairs. The total number of proteins in the test set per tool and the employed performance measures are provided in Table 1. Because no optimal threshold value is reported for these tools, we determine the best threshold per protein using ROC curves for MCC comparisons. We used the same MSA set as an input for all tools in comparison.

**Table 1.** General information about tools and statistical measures employed.

| Tool Name | Input Type | Number of Proteins | Reported Values | Measure Used |
| --- | --- | --- | --- | --- |
| PDB | 3D Structure | 652 | Experimentally Studied Pairs | - |
| PHACE | Phylogenetic Tree, ASR, MSA | 652 | All Position Pairs | AUC, MCC, F1 |
| CoMap | Phylogenetic Tree, ASR, MSA | 652 | Only Co-Evolving Pairs | MCC, F1 |
| CAPS | Phylogenetic Tree, ASR, MSA | 652 | Only Co-Evolving Pairs | MCC, F1 |
| DCA | MSA | 647 | All Position Pairs | AUC, MCC |
| GaussDCA | MSA | 652 | All Position Pairs | AUC, MCC |
| Mlp | MSA | 652 | Missing Values | AUC, MCC |
| PSICOV | MSA | 646 | Missing Values | AUC, MCC |

In line with existing literature, we assessed the performance of PHACE and other tools across diverse scenarios using two distinct test sets focused on co-evolving positions. The first set encompassed all pairs, while the second set specifically included co-evolving pairs with more than 5 amino acids between them. Although the first scenario, comprising co-evolving position pairs with a distance of 5 or fewer amino acids between them, is less extensively detailed in the literature and often perceived as straightforward, our comparison revealed that benchmark tools performed less effectively in this set compared to PHACE. Furthermore, PHACE exhibited significant improvement over these tools even in the second set, which presents more challenging cases. We divided the comparisons into two subsections based on the input of the compared tools.

### Comparison over a Common Set of Proteins

Before diving into detailed pairwise comparisons, we present an AUC-wise comparison of all tools using a common set of proteins comprising 639 entries. It’s crucial to note that for pairs with missing values for any tool in ROC curve comparisons, we assign the respective tool’s lowest score for the corresponding protein. As CAPS and CoMap only report co-evolving position pairs, excluding a common set of positions without a score from any of the six tools was not meaningful.

Fig 3 illustrates the comparison of all tools over two distinct test sets: one constructed over all positional pairs (Fig 3A) and the other over pairs with at least a 5-amino acid separation (Fig 3B). In conducting ROC curve comparisons, we aim to maintain a balanced test set encompassing both co-evolving and independent positional pairs. Independent positions are selected, starting from the furthest pairs to the closest, while minimizing repetitions. Our objective is to maintain a fair comparison and avoid favoring any tool based solely on identical positions. As depicted in Fig 3A and 3B, PHACE demonstrates superior performance compared to all six tools, with a significant difference even compared to the best-performing tool in this set, DCA (t-test, p-value < 0.001).

**Figure 3.**
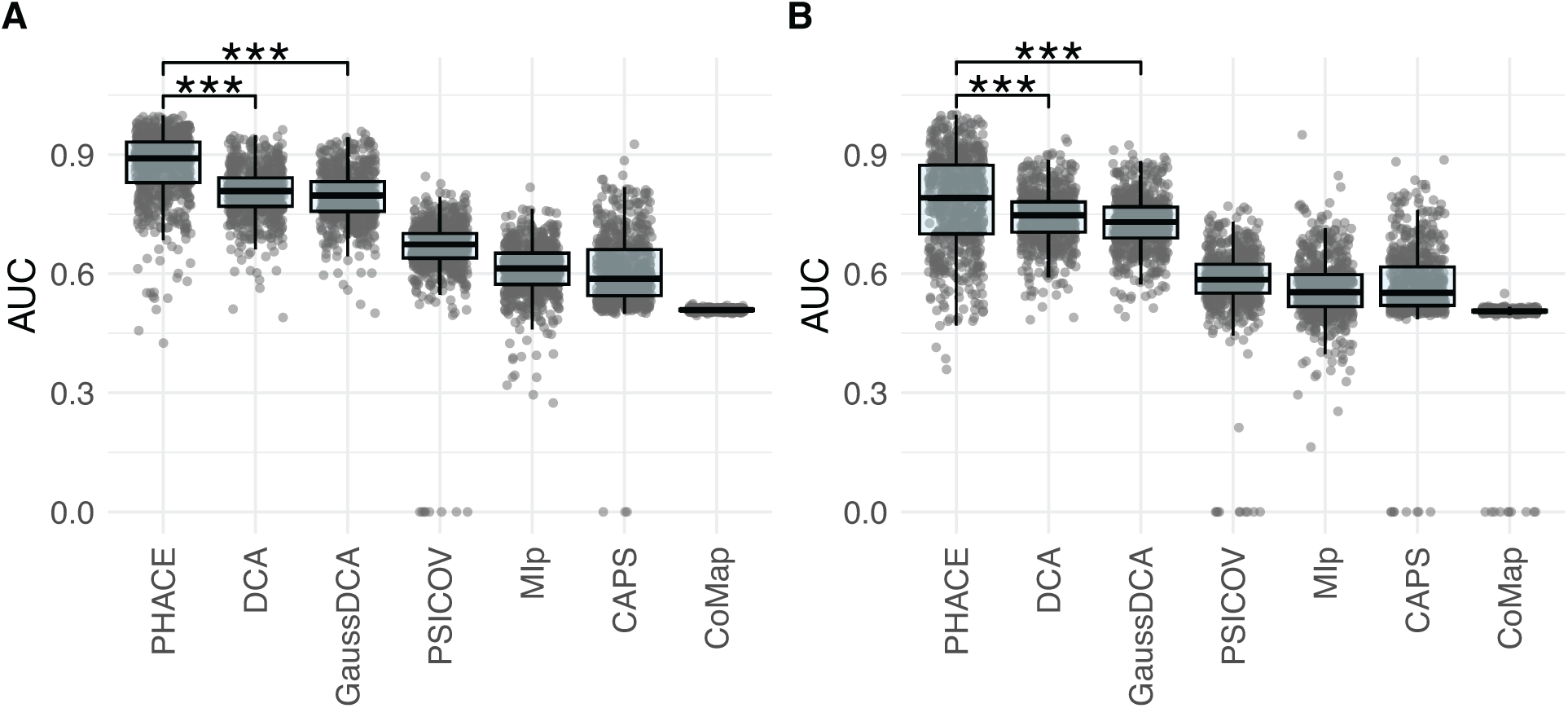
Comparison of all tools over a common set in terms of AUC. The test set is A. all pairs, and B. pairs with at least 5 amino acids between them.

### Comparison among Phylogeny-based Approaches

In the initial series of pairwise comparisons, we assessed PHACE against two other tools, CAPS and CoMap, both of which employ phylogenetic tree analysis and ancestral reconstruction in their predictions. As previously mentioned, CAPS and CoMap specifically identify position pairs considered co-evolving according to their methodologies. CAPS detects coevolving amino acid sites by measuring the correlation of evolutionary rates between sites, adjusted for divergence times. The version we consider (CAPS v2) also incorporates substitution mapping and ASR. However, CAPS has some limitations, such as assigning a single amino acid to internal nodes, ignoring gaps, and relying on a simple correlation function to infer coevolution. Another tree-based tool, CoMap, is a clustering-based method that identifies coevolving amino acid sites by mapping substitutions across a phylogenetic tree. Like CAPS, CoMap assigns a single amino acid to each internal node and represents evolutionary changes in a binary format (0 or 1), potentially oversimplifying the complexity of these changes. Additionally, CoMap also disregards gaps and employs a basic correlation measure. Both approaches do not consider the position dynamics related to tolerable and intolerable amino acids. Since these tools do not generate predictions for all potential pairs, we evaluated them using MCC and F1 scores, which are well-suited for categorical comparison among imbalanced datasets. While a single threshold may not universally optimize performance across proteins with diverse behaviors, we set a threshold (0.25) for PHACE for an equivalent comparison. Fig 4 illustrates the resulting MCC and F1 score performances for pairwise comparisons involving PHACE, CAPS, and CoMap. Fig 4A and 4C correspond to MCC and F1 score comparisons across all possible pairs, while 4B and 4D include pairs with at least a 5-amino acid separation. It is apparent from the figures that PHACE significantly outperforms CAPS and CoMap in terms of both MCC and F1 score for both test sets (t-test p-value < 0.001). This underscores the superior predictive capability of PHACE over these alternative tools that utilize phylogenetic trees in identifying co-evolving position pairs within protein sequences.

**Figure 4.**
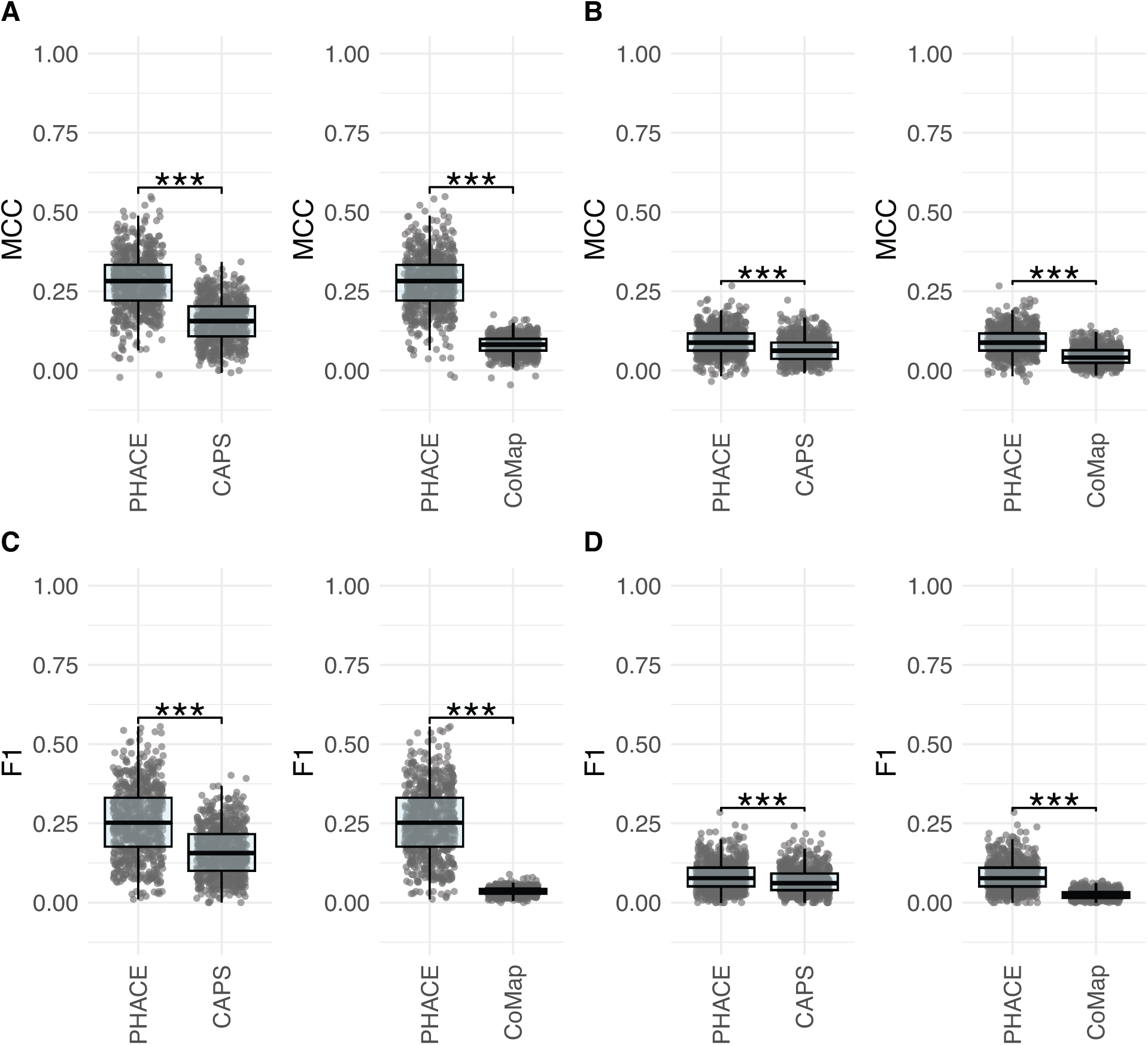
Performance comparison of PHACE and phylogeny-based approaches, CAPS and CoMap in terms of MCC and F1 score. The test set is **A. C.** all pairs, and **B. D.** pairs with at least 5 amino acids between them.

### Comparison of PHACE with MSA-based Approaches

In this section, we comprehensively compared PHACE and several MSA-based tools, namely DCA, GaussDCA, PSICOV, and MIp, focusing on the area under the ROC curve and MCC. We aimed to evaluate the performance of these tools in detecting contacting residues inferred from protein structures.

Similar to the earlier ROC curve comparisons, we aimed to construct a balanced test set comprising co-evolving and independent position pairs. To ensure fairness and avoid favoritism towards any tool based on repeated positions, we selected independent positions starting from the furthest pairs to the closest while minimizing repetitions. The pairs not reported by the compared tool are excluded from the test set. Since each tool may have a different set of test proteins, we conducted pairwise comparisons similar to the previous section. The results in Fig 5 indicate a significant performance gap between PHACE and other MSA-based tools over a test set constructed with all pairs (Fig 5A) and pairs with at least a 5-amino acid separation (Fig 5B). The significance test was again performed using a t-test, with the p-value observed as less than 0.001.

**Figure 5.**
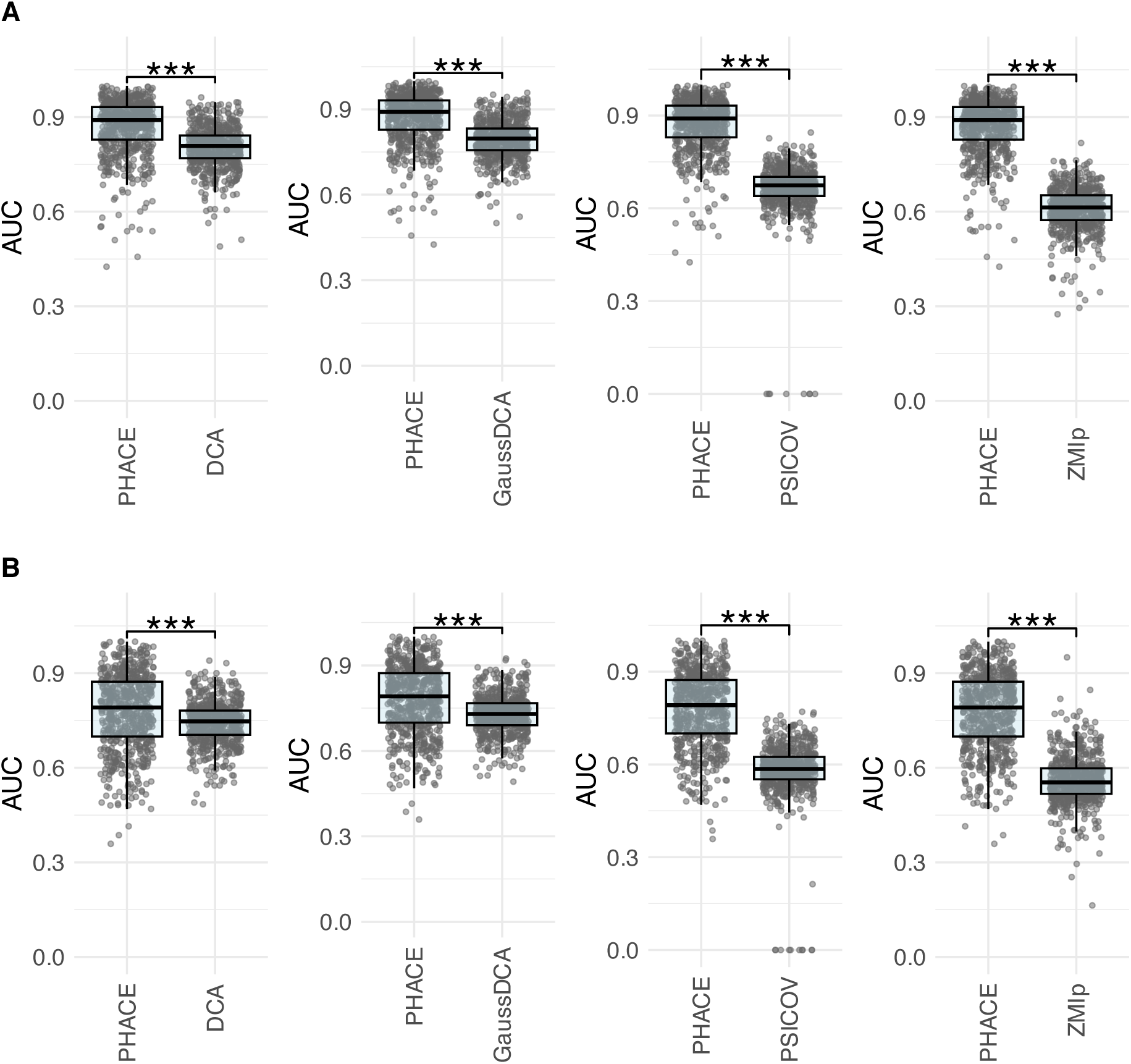
Comparison of PHACE and MSA-based tools, DCA, GaussDCA, PSICOV and MIp in terms of AUC. The test set is constructed over **A.** all pairs, and **B.** pairs with at least 5 amino acids between them.

Transitioning to MCC comparisons, we acknowledged the variability in threshold selection across different tools for individual proteins. To our knowledge, these tools do not report a universally valid threshold. Therefore, we determined the threshold for each tool based on the ROC curve, enabling an unbiased comparison between PHACE and each tool pairwisely in terms of MCC. Fig 6 highlights a statistically significant improvement in PHACE’s performance compared to DCA, GaussDCA, PSICOV, and MIp across test sets over all pairs, as well as pairs with at least a 5-amino acid separation considered. These findings underscore the effectiveness of PHACE in identifying positions in contact within protein sequences, outperforming other established MSA-based tools.

**Figure 6.**
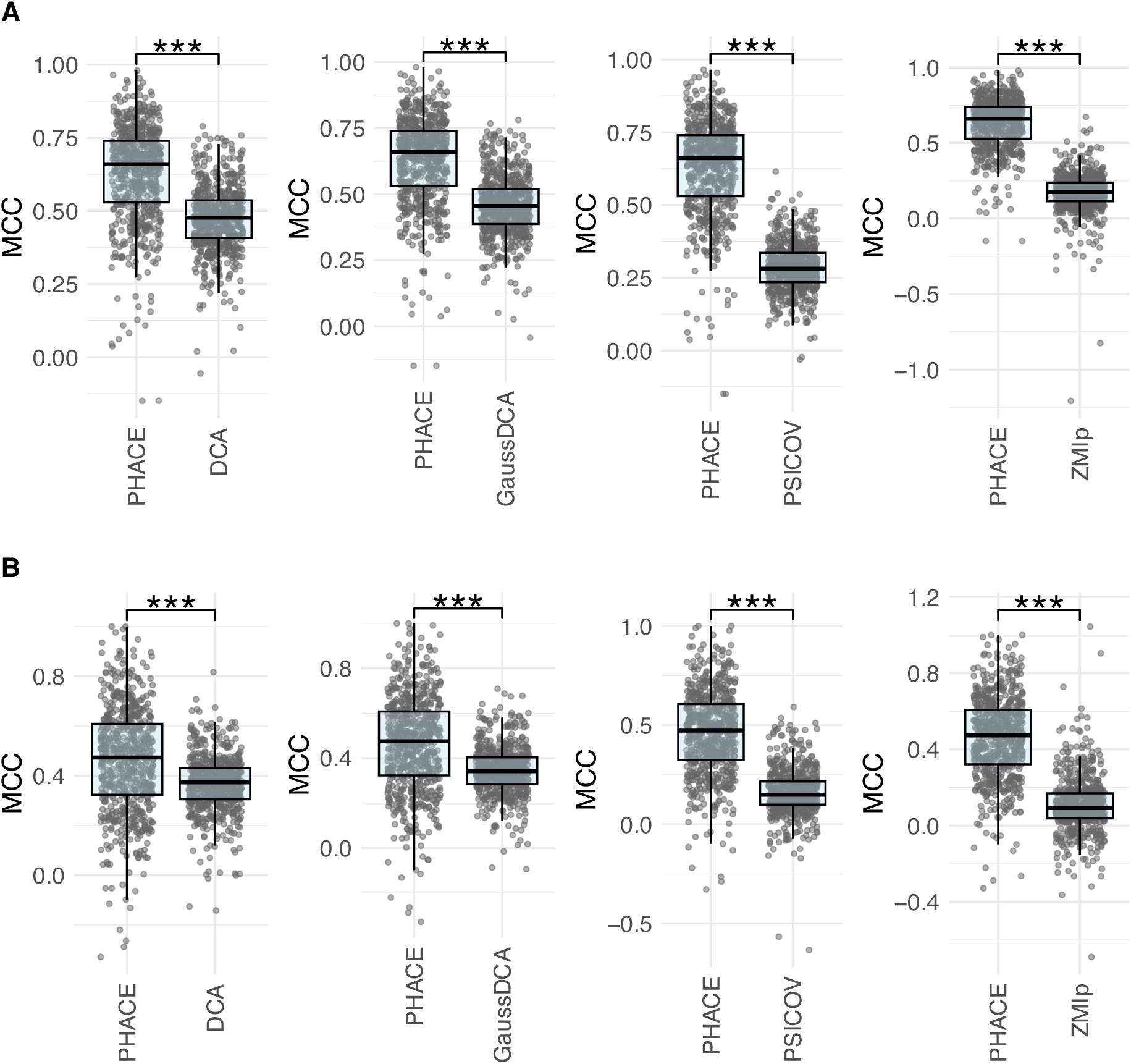
Comparison of PHACE and MSA-based tools, DCA, GaussDCA, PSICOV, and MIp in terms of MCC. The test set is constructed over **A.** all pairs, and **B.** pairs with at least 5 amino acids between them.

### Limitations of Current Approaches

In Fig 7, we aim to illustrate instances where PHACE successfully classifies co-evolving position pairs while other tools fail. These examples shed light on the potential benefits of properly incorporating phylogenetic trees to enhance the prediction of co-evolving positions.

**Figure 7:**
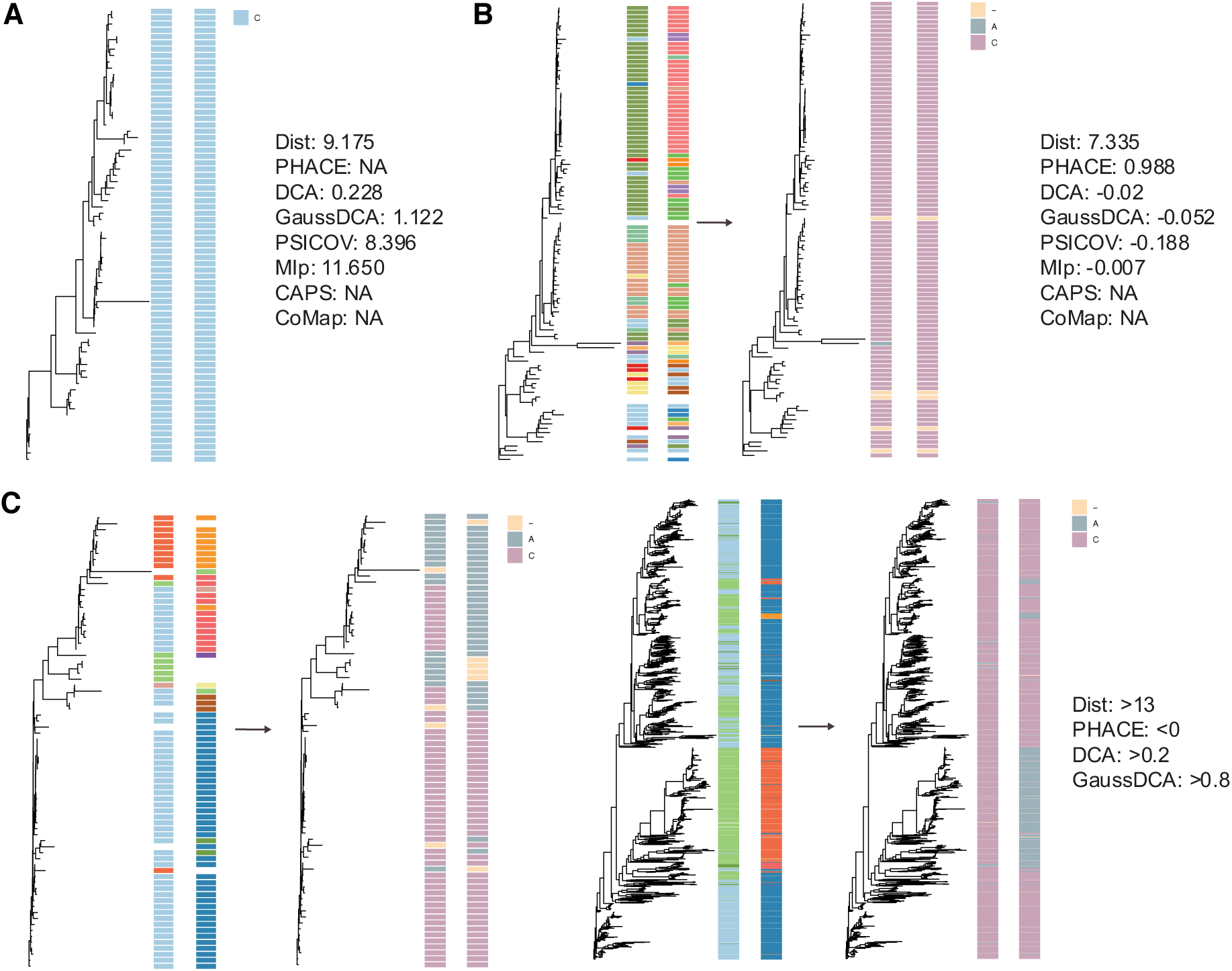
Illustration of selected cases where other tools fail to identify co-evolutionary relationships, while PHACE correctly identifies them. A. Fully conserved position predicted as co-evolving by all MSA-based tools. B. Identification by PHACE when other tools fail. C. Two examples of positions falsely predicted as co-evolving by DCA and GaussDCA.

The first example presents a scenario involving a fully conserved position pair. Despite the absence of any evident signal indicating co-evolution, DCA, GaussDCA, PSICOV, and MIp assign relatively high scores to this pair. We observe instances similar to this, particularly in DCA and GaussDCA. This position pair is strongly predicted as co-evolved by the current tools, although there is no amino acid substitution.

The second example underscores the impact of distinguishing between tolerable and intolerable amino acids based on phylogenetic-independent events. Although the original MSA does not exhibit a strong co-evolution signal for the position pair, the presence of amino acids observed independently during the phylogenetic tree analysis leads to their clustering as tolerable amino acids. Consequently, an updated MSA reveals a noticeable co-evolution signal. As a result, PHACE correctly identifies a pair with a distance of 7.34, while no other tool was able to do so.

The final examples in Fig 7C illustrate the success of tolerable/intolerable clustering in eliminating incorrect co-evolution signals. DCA and GaussDCA predict both pairs as co-evolving with a high score, while PHACE correctly labels them as independent due to phylogenetically independent alterations among the amino acid groups.

These examples highlight PHACE’s ability to effectively leverage phylogenetic information to identify co-evolving position pairs, demonstrating its superiority over other tools in certain scenarios where traditional methods may fall short.

## Discussion

This study introduces a novel perspective on scoring co-evolution among protein positions and presents PHACE, which utilizes phylogenetic trees to assign scores to position pairs based on correlated, phylogenetically independent amino acid alterations. It categorizes observed amino acids into two groups: tolerable amino acids and intolerable amino acids, with gaps considered as the third group of characters. We compared the performance of PHACE with both phylogeny-based approaches (CAPS and CoMap) and state-of-the-art MSA-based tools (DCA, GaussDCA, PSICOV, and MIp). Our results demonstrate a significant difference in performance between PHACE and other benchmark tools across various measures. This improvement is noteworthy as it indicates that by eliminating phylogenetic dependence, a major source of signal that can be mixed with co-evolution, we can achieve better performance than existing state-of-the-art approaches. Moreover, PHACE’s success over phylogeny-based approaches is significant, as while employing phylogenetic trees is crucial to eliminate the correlation of phylogenetic dependency, benefiting from trees to correctly identify phylogenetically independent alterations—the main source of co-evolution—is even more crucial. We believe PHACE achieves this by using a tree traversal process, an approach we have successfully utilized in various problems (Kuru et al. 2022b; Bircan et al. 2023; Dereli et al. 2024). This approach enhances our ability to discern phylogenetically independent alterations accurately, thus contributing to the superior performance of PHACE in identifying co-evolutionary signals among protein positions.

Our analyses evaluated PHACE’s performance using experimentally studied protein structures obtained from the PDB. In line with prevailing literature, position pairs close in 3D structure are commonly assumed to be co-evolving. While this assumption carries the risk of both false positives and false negatives, testing all tools over the same set of co-evolving and independent positions ensures a fair comparison. Our primary objective here is not to predict protein structure. However, leveraging structural data allows us to assess PHACE’s ability to discriminate between co-evolving and independent position pairs based on its scores. We used two thresholds—8 Å and 16 Å— to define co-evolving and non-coevolving pairs for MCC and F1 score comparisons with CAPS and CoMap, respectively. For the ROC curve comparisons with DCA, GaussDCA, PSICOV, and MIp, we implemented a method for selecting non-coevolving position pairs that involved sorting distances from the farthest to the closest up to the 8 Å threshold. This ensured an equal number of co-evolving and non-coevolving pairs to balance the data set, addressing potential biases that could affect the AUC metric, which is more sensitive to imbalances compared to the MCC. This systematic approach maximized the reliability of our comparisons, ensuring that PHACE’s performance was assessed with rigor and consistency across different tools and metrics. By carefully selecting position pairs based on their structural distances, we ensured a comprehensive and fair evaluation of co-evolution predictions across different assessment methods.

In our comparisons, we employed two distinct but overlapping test sets. The first set encompassed all position pairs, while the second set comprised position pairs with at least five amino acids between them. The rationale for this division is rooted in the literature, which suggests that the second set presents more challenging cases, as pairs with fewer than five amino acids between them are considered easier to predict. However, our observations deviate from these expectations. While there was a slight performance increase for all tools considered, none of the six tools achieved fully successful predictions. Moreover, the performance gap between PHACE and all six tools widened when considering the test set encompassing all pairs, including the “easy” ones.

We want to highlight that we disregarded the unreported proteins and conducted pairwise comparisons exclusively over each tool’s reported set of proteins. Furthermore, even when results were obtained for a protein, DCA, GaussDCA, PSICOV, and MIp might not have produced results for all possible position pairs. In such cases, comparisons were made by excluding the unreported pairs, potentially providing an advantage for the tool in question. On the other hand, PHACE successfully reports a score for all position pairs within a protein. Therefore, we did not exclude any protein or position pair based on PHACE predictions. Although excluding proteins/positions based on a tool’s prediction might have brought an advantage for that tool, PHACE consistently outperformed these tools in the comparisons.

Fig 6 visually demonstrates PHACE’s superiority over other benchmark tools. Particularly noteworthy is our clustering approach, which considers the tolerance of positions to amino acid alterations, resulting in a notable performance enhancement compared to other tools. It’s worth mentioning that DCA, GaussDCA, PSICOV, and MIp may assign a high score, indicating co-evolution for conserved position pairs. However, we excluded these pairs from our comparisons as they deviate from the definition of co-evolution, which entails correlated changes between positions.

Motivated by the substantial performance enhancement achieved with PHACE, our next step is to extend our approach to detect protein-protein interactions. Protein-protein interactions play a pivotal role in various cellular functions, and it’s well-established that many human diseases arise from abnormal protein–protein interactions (Ryan and Matthews 2005). However, detecting these interactions through experimental methods is time-consuming and expensive (Macalino et al. 2018; Chen et al. 2019) while current computational approaches have yet to reach the desired accuracy level (Gandarilla-Pérez et al. 2023). One potential avenue for improving the prediction of protein-protein interactions is to generate enhanced co-multiple sequence alignments (co-MSAs), where each row represents a combination of two interacting proteins. Our initial objective is to develop a phylogeny-aware algorithm to construct reliable co-MSAs. Subsequently, PHACE might be useful in predicting protein-protein interactions.

As another future direction, we aim to enhance PHACT by integrating co-evolution information obtained from PHACE scores. PHACT predicts the pathogenicity of missense mutations by utilizing phylogenetic trees and phylogenetically independent amino acid alterations. While it is an accurate variant effect predictor, PHACT currently assumes each protein position to be independent, which is an incorrect assumption. It would be useful to incorporate co-evolution information and the branches contributing to co-evolution into the PHACT algorithm to improve its performance.

## Materials and Methods

### Details of PHACE

PHACE comprises three crucial components in its algorithm design.

#### 1. Constructing MSA_1_

Initially, we detect tolerable/intolerable amino acids by determining the amino acid with the highest frequency at each corresponding position in the MSA. This amino acid serves as a baseline for identifying tolerable amino acids.

Tolerable and intolerable amino acids are determined based on their scores computed over phylogenetically independent substitutions. We traverse the tree from the root, assessing the probability difference per amino acid over neighboring nodes. The final score is derived through weighted summation of positive probability differences. Amino acids with scores higher than the baseline are labeled tolerable; otherwise, they are considered intolerable. In the first alternative MSA, MSA_1_, we designate the character “C” for tolerable amino acids, “A” for intolerable amino acids, and maintain gaps as they are.

To compute the total phylogenetically independent change per branch, we traverse the tree, calculating the summation of positive probability differences per branch. Thus, we have a matrix of number of branches by 2 including total change per branch.

### 2. Constructing MSA_2_

The limitation with MSA_1_ and the total changes computed over MSA_1_ is that the gap character is not considered in the ASR step. Consequently, the probability distribution is focused solely on characters A and C, disregarding gaps. This oversight poses an issue, as branches where the probability of a character increases may erroneously include substitutions to gaps, even if those gaps did not occur phylogenetically independently. To address this issue, we introduce a second MSA, MSA_2_, comprising two characters: “C,” representing all 20 amino acids, and “G” for gaps. With MSA_2_, we rerun ASR and apply the same tree traversal process as with MSA_1_. This enables us to identify branches where phylogenetically independent substitutions to G occur, along with the corresponding amount of change.

We then update the initial matrix constructed over MSA_1_ with information regarding the branches where gap alterations occur and the associated amount of change. This update ensures that our matrix encompasses all phylogenetically independent alterations, thereby providing insights into co-evolution through correlation analysis.

### 3. Score Computation

The weighted concordance correlation coefficient (WCCC) serves as a pivotal metric in our analysis, particularly for quantifying the parallelity between the total amount of changes for branches per position. While traditionally employed to measure agreement between two variables, WCCC proves invaluable in our context due to its ability to assess correlation while accounting for both the magnitude of change and the importance of each branch through the application of weights.

To adapt WCCC to our specific needs, we’ve refined the original formula to incorporate these considerations. The updated formula is as follows:

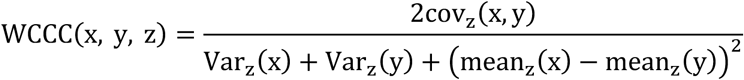

where x and y represents the total amount of change per branch for position 1 and 2 in the pair, respectively and the subscript z corresponds to the weighted version, where each term is weighted by the weight associated with the branch. This refined formulation of WCCC enables us to effectively capture the nuanced relationship between changes across branches and positions, while accommodating variations in the importance of individual branches in terms of co-evolution signal. Thus, it serves as the most suitable measure for our analytical needs.

We utilize two distinct weights in PHACE: one pertains to the incompatibility related to gap characters, denoted as ω_1_, while the other is assigned per branch. The formula of the first weight is as follows

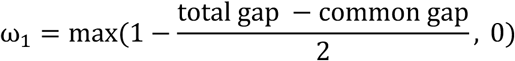

total gap refers to the total number of branches with gaps for the first and second positions in the pair. common gap corresponds to the number of branches with gaps that are common for both positions.

The second weight, ω_”_, reflects the diversity of each branch in terms of phylogenetically independent alterations across all positions. However, to ensure that each branch contributes proportionately to the final score relative to the amount of change, we take the geometric mean of branch diversity and the maximum amount of change per branch over the position pair. The formula for the weight per branch i is as follows

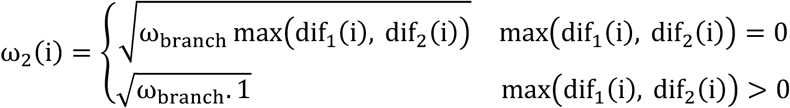

where ω_branch_ is the weight computed over branch diversity and dif_1_(i) and dif_”_(i) correspond to the total change for branch i for the first and second positions in the pair, respectively. We note that if there is a non-parallel change (|dif_1_(i) − dif_”_(i)| ≥ 0.5) on branch i, we assign ω_”_(i) = 1 to ensure that the effect of non-parallel change is not reduced.

The final PHACE score is computed by considering both weights and WCCC as follows

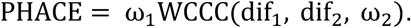

Here, it’s important to note that in the case of a non-parallel change, we examine the original MSA. If the amino acid in question is observed only once, we disregard the impact of this change and assume that there is no change on the corresponding branch for both positions in the pair. Additionally, substitutions between amino acids and substitutions to gaps are not considered correlated changes, even if they occur on the same branch for both positions. We penalize the score for these types of parallel changes.

### PDB Structures

The experimentally studied protein structures are acquired using a batch download script directly from the PDB (Berman et al. 2002). For each UniProt ID, the corresponding PDB ID is retrieved from the UniProt database (UniProt 2021). For 2,390 proteins which we have from (Kuru et al. 2022a), have PDB structures available. To assess the compatibility of the sequences in the structures, we collected three information: number of compatible positions, number of different positions and if the sequence at PDB is longer, the length difference between our sequence and PDB sequence. If a structure has more than 10 incompatible amino acid positions or if the ratio of mapped to the total sequence length is less than 50%, it is discarded. From the remaining proteins and structures, if there are multiple candidate structures for a protein, we select the one with the highest number of compatible and minimum number of incompatible positions. That resulted in 652 proteins in total.

### Benchmark Tools

We utilized CAPS, CoMap, DCA, GaussDCA, PSICOV, and MIp as benchmark tools, obtained from the GitHub page or web server of the corresponding tool. For a completely fair comparison, each tool was executed over the masked MSA and phylogenetic tree, if required, which are also used for PHACE computation.

The details regarding the parameters are as follows:

- CAPS was executed with default parameters.
- The “Correlation” version of CoMap was employed.
- DCA and GaussDCA were run with default parameters over the masked MSA, except GaussDCA, which reported position pairs with at least 5 amino acids between them. To obtain their predictions over all pairs, we changed the parameter min_separation to 1.
- PSICOV was run with the minimum sequence separation parameter set to 1, similar to GaussDCA.
- MIp was executed with default parameters.

### Multiple Sequence Alignment & Phylogenetic Trees

The MSA and phylogenetic trees of 5,123 human proteins are obtained from the PHACT database (Kuru et al. 2022a). They obtained the homologues of each query sequence through PSI-BLAST (Altschul et al. 1997) against a non-redundant database of 14.010.480 proteins produced from the reference proteomes in the UniProtKB/Swiss-Prot Knowledgebase (UniProt 2021). Two iterations of PSI-BLAST with 5000 maximum target sequences were performed. The number of hits were limited to maximum 1000 sequences with a minimum 30% identity and E-value of 0.00001 due to computational limitations of building phylogenetic trees. The sequences were aligned using MAFFT FFTNS (Katoh and Standley 2013), and the multiple sequence alignments (MSAs) were trimmed with the trimAl tool gappyout method (Capella-Gutierrez et al. 2009). The resulting MSA was used to generate a maximum-likelihood phylogenetic tree with the RaxML-NG (Kozlov et al. 2019) tool using LG4X model and leaving the remaining parameters at default settings.

### Ancestral Reconstruction

Positions with ‘gap’ character in the query sequence are removed from the original MSA (without trimming). The resulting MSA is used to perform ancestral sequence reconstructions by using IQTREE. To ensure that amino acid properties do not influence the resulting probability distributions, we employed a user-defined model that assigns equal substitution rates and baseline frequencies to each character. ASR is executed for three versions of the multiple sequence alignment (MSA):

i) The original MSA used to compute tolerance scores per position.
ii) MSA with three characters: the dominating amino acid, the alternating amino acid, and gaps.
iii) MSA with two characters: one character representing all amino acids and another representing gaps.

A similar user-defined model is applied to all three versions, with matrix sizes adjusted based on the number of characters in the MSA. While the tree topology is preserved in the ASR step, it re-optimizes the branch length. To prevent changes in branch lengths based on alternative MSAs, we utilize the -blfix option, which ensures fixed branch lengths.

## Acknowledgements

This study is supported by the Scientific and Technological Research Council of Turkey (TÜBİTAK). The numerical calculations reported in this paper were performed at TOSUN cluster at Sabanci University and TUBITAK - Turkish Academic Network and Information Center (ULAKBIM), High Performance and Grid Computing Center (TRUBA resources).

## Data Availability

All data generated in this study and all benchmark analysis scripts and source codes for PHACE are available at https://github.com/CompGenomeLab/PHACE. The PHACE predictions for the 652 proteins used in this manuscript are provided at https://zenodo.org/records/14043199.

